# Integration of behavioral and biological variables using penalized regression: an application to the maternal immune activation model of autism

**DOI:** 10.1101/2020.03.31.018333

**Authors:** Cristina Paraschivescu, Susana Barbosa, Thomas Lorivel, Nicolas Glaichenhaus, Laetitia Davidovic

## Abstract

Maternal immune activation (MIA) during pregnancy increases the odds of developing neuropsychiatric disorders such as autism spectrum disorder (ASD) later in life. In pregnant mice, MIA can be induced by injecting the viral mimic polyinosinic:polycytidylic acid (poly(I:C) to pregnant dams resulting in altered fetal neurodevelopmental and behavioral changes in their progeny. Although the murine MIA model has been extensively studied worldwide, the underlying mechanisms have only been partially elucidated. Furthermore, the murine MIA model suffers from lack of reproducibility, at least in part because it is highly influenced by subtle changes in environmental conditions. In human studies, multivariable (MV) statistical analysis is widely used to control for covariates including sex, age, exposure to environmental factors and many others. We reasoned that animal studies in general, and studies on the MIA model in particular, could therefore benefit from MV analyzes to account for complex phenotype interactions and high inter-individual variability. Here, we used a dataset consisting of 26 variables collected on 67 male pups during the course of several independent experiments on the MIA model. We then analyzed this dataset using penalized regression to identify variables associated with *in utero* exposure to MIA. In addition to confirming the association between some previously described biological variables and MIA, we identified new variables that could play a role in neurodevelopment alterations. Aside from providing new insights into variable interactions in the MIA model, this study highlights the importance of extending the use of MV statistics to animal studies.

## 1. Introduction

Maternal immune activation (MIA) increases the child’s odds of developing neuropsychiatric disorders later in life [1]. Notably, *in utero* fetus exposure to maternal infection is a risk factor for developing schizophrenia (SCZ) and autism spectrum disorders (ASD) [1-4]. MIA can be modelled in rodents by gestational administration of the viral mimic poly(I:C) [3, 5, 6], which triggers a broad inflammatory response accompanied by heightened levels of peripheral maternal proinflammatory cytokines such as interleukin (IL)-6 and IL-17A [7-9]. Exposure to MIA triggers irreversible neurodevelopmental defects in the fetus, eventually resulting in behavioral alterations in the socioemotional domain frequently observed in patients with ASD or SCZ [5].

In most human studies, MV statistical analysis is used to control for multiple covariates including sex, age, exposure to environmental factors and many others. This is not the case in the vast majority of animal studies in which experimental conditions are well controlled and in which inter-individual variability is reduced. Hence, in animal studies in which complex phenotypes are analyzed and/or inter-individual variability is difficult to control, standard parametric or non-parametric analysis between groups (Student’s t-test, Mann-Whitney test, ANOVA, Kruskal-Wallis) or pairwise correlations between variables (Pearson’s correlation, Spearman’s correlation) only provide limited insight into the complexity of the interactions between variables. This is particularly the case in the MIA model in which the progeny phenotype can be heavily influenced in unpredictable ways by experimental conditions [10, 11]. We therefore believe that animal studies in general, and notably the MIA model, could benefit from MV analyzes to model the effects of variables on the outcome of interest.

Several MV methods are currently available including principal component analysis (PCA), and linear and logistic regressions. PCA is an unsupervised method, often used to detect groups of related samples. However, PCA does not allow for identifying the variables responsible for group separation, and its discriminatory potential is limited as variables that contribute little towards explaining variation between groups are not eliminated. These issues can be overcome by supervised approaches, such as logistic regressions, in which the model is informed of the sample’s class (e.g., untreated vs. treated) therefore maximizing inter-class discrimination. Among different regression methods, the penalized regression framework is well adapted to datasets in which the number of events per variable is low (in other terms when the sample size is small and the number of variables under scrutiny is high) and /or there is multi-collinearity among variables (for example when variables are highly correlated) [12], as frequently observed in datasets from animal studies.

Here, we used a dataset consisting of 26 variables collected on 67 male pups during the course of several independent experiments on the MIA model. We then analyzed this dataset using a penalized regression approach to identify variables associated with *in utero* exposure to MIA.

## 2. Materials and Methods

### 2.1. Animals

Animal housing and experimentation were conducted in facilities certified by regional authorities (Direction Départementale de Protection des Populations des Alpes-Maritimes, accreditation #C-06-152-5). The study was in accordance to procedures approved by the Ministère de l’Enseignement Supérieur et de la Recherche and the local ethics committee for animal experimentation (Ciepal Azur), in agreement with the EU Directive 2010/63/EU for animal experiments. Given the recent demonstration that C57Bl/6N Taconic mice were more susceptible to MIA-induced behavioral defects [9], twelve female and four male C57Bl/6N founder mice were purchased from Taconic Biosciences (Lille Skensved, Denmark). Mice were housed in open medium cages equipped with wooden litter, cotton to nidify as well as a one plastic house, in a temperature (22-24°C) and hygrometry (70-80%)-controlled room, with a 12-h light/dark cycle (lights on from 8:00 a.m. to 8:00 p.m.) with food (standard chow, reference 4RF25, Mucedola) and water *ad libitum*. Mice were housed by 3-5 animals per cage.

### 2.2. Experimental procedures

To obtain a high-confidence dataset, we followed the recommended guidelines for the MIA model [10, 11]. Females were mated with males for 16h (6 p.m.-10 a.m. next day, considered embryonic day (E) 0.5), by groups of 3 females for 1 male. After mating, females were left undisturbed, with the exception of weekly cage cleaning. Pregnant dams (identified based on minimal body weight (BW) gain of 3 g between E0.5-E11.5) were randomly assigned to the experimental groups and injected with poly(I:C) (n=13) or vehicle (saline, n=8) at E12.5 between 9-10 p.m. Due to previously reported intra-lot variability of poly(I:C) [11, 13], a single lot (reference P9582, Sigma-Aldrich) was dissolved in sterile double-distilled water at room temperature (RT) to generate a stock solution based on pure poly(I:C) weight (20µg/µL). Stock solution was aliquoted and stored at −20°C until use. For each cohort, a new aliquote was diluted in 0.9% NaCl to the working concentration of 1µg/µL. Pregnant dams received a single intraperitoneal injection of either poly(I:C) at a dose of 5 mg/kg (5µL/g body weight of the 1µg/µL solution) or 5µL/g body weight of saline.

### 2.3. Collected variables

#### 2.3.1. Biological variables

Maternal and paternal ages were recorded. Maternal rectal temperature was recorded before, as well as 3 and 6 h post injection. Body weight (BW) was recorded before and 24 h post injection. At E16.5, the pregnant dams were separated into individual cages until birth, considered as postnatal day 0 (P0) and left undisturbed until P4. Litter size was recorded at P4. Only males were included in the study, but females remained in the litter to preserve numbers and sex balance. Male pups were individually marked at P4. We followed a total of n = 27 male pups born from mothers injected with saline (from 8 litters spread across 3 independent cohorts) and n = 40 male pups born from mothers injected with poly(I:C) (from 12 litters spread across 4 independent cohorts).

At P15, BW was recorded and pups were euthanized using a lethal dose of anesthetic. Blood was collected by cardiac puncture into 1.5 ml Eppendorf tubes, allowed to clot for 30 min RT and centrifuged (10,000g, 4°C, 20 min) to recover serum stored at −80°C. The V-PLEX® Mouse Cytokine 29-Plex Kit electrochemiluminescence-based assays from Meso Scale Diagnostics (Rockville, USA) was used to measure IFN-γ, IL-1β, IL-2, IL-4, IL-5, IL-6, IL-12p70, CXCL1, TNF-α, IL-9, CCL2, IL-33, IL-27p28/IL-30, IL-15, IL-17A/F, CCL3, CXCL10, CXCL2, CCL20, IL-22, IL-23, IL-17C; IL-31, IL-21, IL-16, IL-17A, IL-17E/IL-25. Assays were performed according to the manufacturer’s instructions. Cytokines (IL-2, IL-4, IL-12p70, IL-9, IL-17A/F, IL-22, IL-23, IL-17C, IL-31, IL-21, IL-17E, IL-25) below the Lower Limit of Detection (LLOD) in more than 20% of the samples were excluded. For the other cytokines, cytokine levels below LLOD were imputed a value equal to half the LLOD value indicated by the manufacturer.

#### 2.3.2. Behavioral variables

At P6, individual pups were isolated from their mother and litter and placed on a cotton-padded dish into a thermo-controlled (26°C) soundproof chamber. Ultra-sonic vocalizations (USV) were recorded for 5 min using the UltraSoundGate Condenser Microphone and 116 USB Audio device (Avisoft Bioacoustics), as described [14, 15]. Four pups were recorded simultaneously in parallel chambers. Sonograms were analyzed with the AvisoftSASLab Pro software (version 5.2.12, Avisoft Bioacoustics) based on automated recognition of USV using a 25 kHz cut-off frequency and a 2-7 ms element separation, as described in [14]. Misidentified USV were manually curated and number and duration of USV were automatically extracted. At P13, pup was placed in a clean individual cage and its movements recorded during 10 min with a digital camera. Videos were analyzed using the ANY-maze video-tracking software (Dublin, Ireland), which extracted the total distance travelled and the time spent mobile for each individual.

### 2.4. Statistics

#### 2.4.1. Correlation studies

For correlation studies, pairwise correlations between the 25 numerical variables (all variables except multiparity) and associated p-values were assessed using the Spearman’s r correlation coefficient rank test. We did not adjust the p-values for multiple testing as this correlation analysis was purely descriptive to highlight possible relationship between variables (multi-collinearity) and not meant for biological interpretation.

#### 2.4.2. Penalized regression

##### Rationale for the choice of penalized regression to analyze datasets derived from animal studies

Regression problems with many potential candidate and adjustment variables usually require model selection to find a simpler model, while keeping a good performance. While penalized regression methods are mostly used in high-dimensional settings, their usefulness in low-dimensional has been shown [16]. Indeed, traditional stepwise selection methods, such as forward and backward selection, suffer from high variability and low accuracy, in particular in settings with large number of covariables or when there is correlation between covariables [17]. Penalized methods, such as LASSO, Elastic Net and Ridge regressions are an alternative to such traditional methods. Alike ordinary least squares (OLS) estimation, these methods estimate the regression coefficients by minimizing the residual sum of squares (RSS), while placing a constraint on the size of the regression coefficients. This constraint or penalty has the disadvantage of biasing the coefficient estimates, however, it improves the overall prediction error of the model by decreasing the variance of the coefficient estimates or odd-ratios (OR).

A penalized regression method yields a sequence of models, each associated with specific values of the α and λ hyperparameters. α accounts for the relative importance of the L1 (LASSO) and L2 (ridge) regularizations and λ controls the magnitude of regularization [18]. Thus, a tuning method for one or both hyperparameters needs to be specified to achieve an optimal model. There are several tuning methods such as AIC, Cp statistic, average square error on the validation data and cross-validation. By minimizing the RSS using a penalty on the size of the regression coefficients, some regression coefficients will shrink towards zero. If the penalty is extreme, regression coefficients are set to zero exactly. Thus, penalized regression performs both variable selection and coefficients regularization, enhancing the prediction accuracy and interpretability of the resulting statistical models [19].

##### Implementing penalized regression

###### Building the dataset

All variables collected for each individual pup involved in the study were gathered in a matrix (n=67 lines corresponding to the 67 pups; p=26 column corresponding to the 26 variables collected). Penalized regression does not allow for missing values. In this study, we only considered individuals for which all variables were available. In some cases, to avoid discarding individual samples presenting missing values, imputation of missing values can be achieved using for example the Multiple Imputation by Chained Equations (MICE) package in R [20]. As recommended, for a fair comparison of the relative predictor importance across all explanatory variables, all numerical variables were mean-centered and then scaled such that the input matrix has all column-means equal to 0 and column-variances equal to 1 [21]. Categorical variables were encoded as numbers, with each category represented with a binary vector that contains 1 and 0, denoting the inclusion in one category or the other. The number of vectors depends on the number of levels for the categorical variable and full encoding means all levels are included in the analysis. In our case, the categorical variables considered were binary: multiparity (No/Yes) and the outcome (Control/MIA).

###### Resampling and penalized regression

Fig 1 depicts the workflow of the analysis.

**Fig 1.**
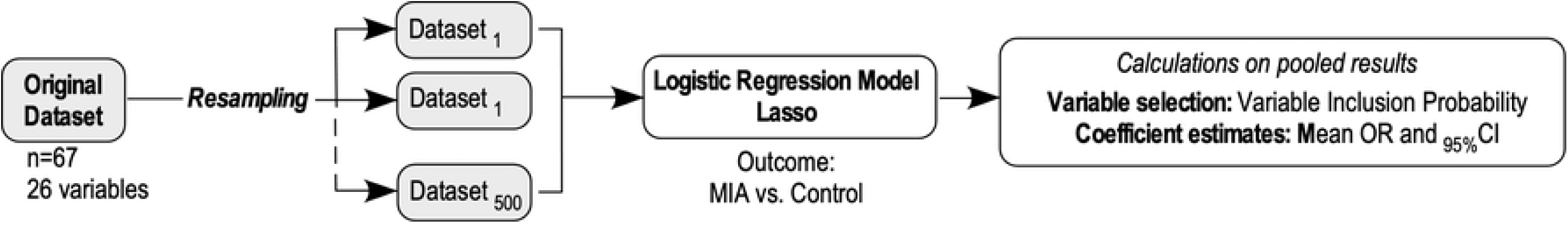
Methodological workflow for the multivariable analysis.

To model the association between parental and pups variables and the outcome “belonging to the MIA class”, we implemented a penalized regression (Lasso model) using the glmnet and caret R packages [18, 22]. Of note other softwares are available (e.g. PROC GLMSELECT in SAS). Considering that current penalized regression methods do not provide valid confidence intervals, or p-values, for testing the significance of coefficients, we used the non-parametric bootstrap for inference [21]. The bootstrap step involved 500 resamplings of the dataset to create 500 different samples of the same size. For each of the resampling runs, we obtained the list of the variables selected by Lasso logistic regression model and the corresponding OR. Based on these, we computed for each variable: 1) the variable inclusion probability (VIP), as the percentage of the 500 bootstrap resamples in which each variable was selected by the penalized regression; 2) the mean OR, computed across the 500 bootstrap resamples, and 3) the non-parametric confidence intervals (CI). The CI was determined by first ordering the bootstrap penalized coefficient estimates from the lowest to highest, then selecting values at the chosen percentile for the confidence interval. For a confidence interval of 95%, as the lower bound we selected the OR value at the 2.5% percentile and as the upper bound of the CI the 97.5% percentile [23].

###### Interpretation of VIP and coefficients

We used the VIP as a measure of the stability of the association between the variable of interest and the outcome. Depending on the study, determining an appropriate threshold for the VIP can be challenging. In their seminal paper, Bunea et al. 2011 recommended to use a “conservative threshold of 50%” because their goal was “not to miss any possibly relevant predictors” [21]. However, this 50% threshold increases the risk of false positives. In this study, we chose to use a stringent VIP threshold of 80% to decide whether the variable under scrutiny was associated with the outcome.

Then, we used the mean OR for the selected variables as a measure of the strength of the association between the variable and the outcome. The OR is calculated as the exponential function of each logistic regression coefficient. The OR represents the odds that an outcome will occur given a particular variable, compared to the odds of the outcome occurring in the absence of that variable [24]. In this study, we used the mean OR to determine whether a particular variable is a risk factor for belonging to the MIA class, and to compare the magnitude of various variables for that outcome. In terms of interpretation, a variable with an OR>1 is associated with higher odds of belonging to the MIA class and a variable with OR<1 is associated with lower odds of belonging to the MIA class.

Finally, the 95% CI was used to estimate the precision of the OR. A large CI indicates a low level of precision of the OR, whereas a narrow CI indicates a higher precision of the OR.

## 3. Results

Our dataset consisted of 26 variables collected longitudinally on 67 male pups (MIA, n=40; control, n=27): paternal and maternal age, multiparity, maternal hypothermia and body weight loss (a surrogate of the intensity of MIA), litter size, pup’s body weight, pup’s behavioral (time spent mobile and distance travelled, number of emitted USVs) and biological (serum levels of IFN-γ, IL-1β, IL-5, IL-6, CXCL1, TNF-α, CCL2, IL-33, IL-27p28/IL-30, IL-15, CCL3, CXCL10, CXCL2, CCL20, IL-16, IL-17A) variables. We first investigated the correlation (multi-collinearity) between the 25 numerical variables of the dataset, using the Spearman’s r correlation coefficient rank test (Fig 2). Data showed medium-to-high correlations among several variables therefore providing a rationale for the use of penalized regression.

**Fig 2.**
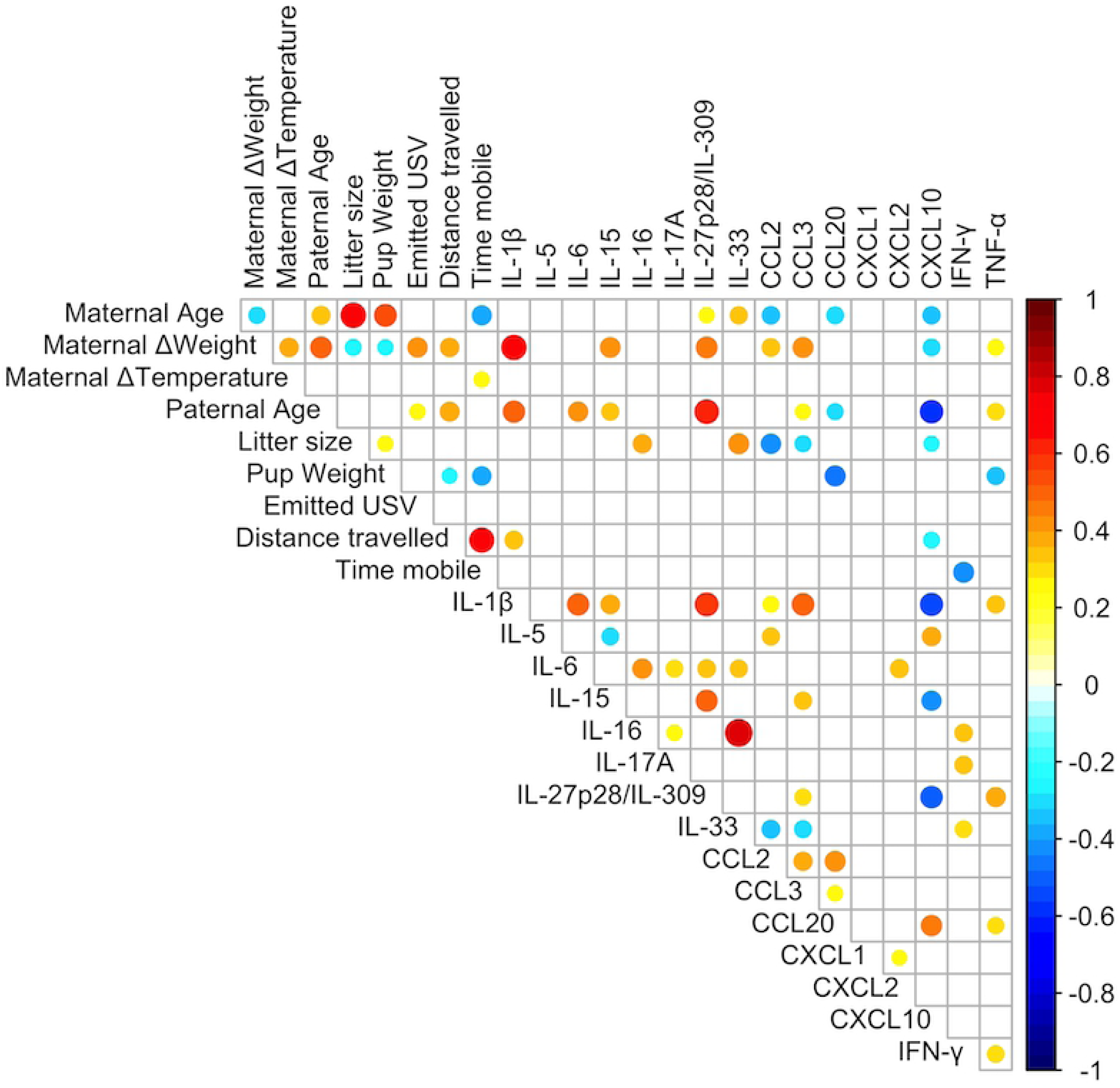
Correlation analysis for the 25 numerical variables present in the MIA dataset. Heatmap of the pairwise Spearman’s rank r correlation coefficient between all variable pairs. Spearman’s r coefficients are color-coded and proportional to dot area. Only significant correlations are displayed (p < 0.05).

Analyzing the dataset using penalized regression identified several variables associated with higher odds of belonging to the MIA class: decrease in maternal body temperature and weight loss, smaller litter size, increased pup weight, reduced number of emitted USVs, reduced distance travelled and reduced time spent mobile, higher serum levels of IL-15 and TNF-α, and lower serum levels of CXCL10 and IL-5 (Table 1).

**Table 1.** Adjusted associations between parental, pup’s variables and cytokine levels and belonging to the MIA class. Variable Inclusion Probability (VIP), mean Odd Ratios (OR) and percentile bootstrap 95% Confidence Interval (CI) computed from penalized regression runs are shown for each of the variables. The VIP is used as a measure of the stability of an association, as it can be interpreted as the posterior probability of including a given variable in the model. Variables with VIPs above 80% (shaded in grey) were considered as stably associated with the MIA class. In the OR column, positive (shaded red) and negative (shaded blue) associations are indicated

|  | Stage | Variable description | VIP | OR | 95% CI |
| --- | --- | --- | --- | --- | --- |
| **Parental variables** | E0.5 | Maternal age (weeks) | 65.7% | 1.139 | [1.009,1.411] |
|  | E0.5 | Multiparity (Yes) | 17.2% | 1.215 | [1.009,2.098] |
|  | E13.5 | **Maternal Δweight (g)** | **99.4%** | **0.044** | **[0.001,0.514]** |
|  | E12.5 | **Maternal Δtemperature (°C)** | **92.1%** | **0.217** | **[0.032,0.826]** |
|  | E0.5 | Paternal age (weeks) | 57.3% | 1.03 | [0.928,1.197] |
| **Pups' variables** | P4 | **Litter size (# pups)** | **93.0%** | **0.334** | **[0.061,0.898]** |
|  | P15 | **Body weight (g)** | **98.7%** | **7.568** | **[1.481,127.402]** |
|  | P8 | **Emitted USV (#)** | **97.0%** | **0.992** | **[0.983,0.998]** |
|  | P13 | **Distance travelled (m)** | **82.6%** | **0.523** | **[0.183,0.956]** |
|  | P13 | **Time mobile (s)** | **90.8%** | **0.989** | **[0.972,0.999]** |
| **Cytokines** | P15 | CCL2 | 57.4% | 0.848 | [0.48,1.25] |
|  | P15 | CCL3 | 63.9% | 0.704 | [0.283,1.065] |
|  | P15 | CCL20 | 43.9% | 1.007 | [1,1.04] |
|  | P15 | CXCL1 | 73.2% | 0.97 | [0.9,1.014] |
|  | P15 | CXCL2 | 70.5% | 0.867 | [0.501,1.096] |
|  | P15 | **CXCL10** | **91.5%** | **0.956** | **[0.881,0.995]** |
| | P15 | IFN-γ | 59.8% | 0.786 | [0.069,86.478] |
| | P15 | IL-1β | 57.2% | 1.338 | [0.807,2.88] |
|  | P15 | **IL-5** | **85.6%** | **0.823** | **[0.588,0.993]** |
|  | P15 | IL-6 | 58.1% | 0.965 | [0.831,1.134] |
|  | P15 | **IL-15** | **85.6%** | **1.035** | **[0.998,1.118]** |
|  | P15 | IL-16 | 36.3% | 1 | [1,1] |
|  | P15 | IL-17A | 53.4% | 0.866 | [0.007,107.497] |
|  | P15 | IL-27p28/IL-309 | 74.7% | 1.344 | [0.975,2.32] |
|  | P15 | IL-33 | 58.0% | 1.052 | [1.002,1.175] |
|  | P15 | **TNF-α** | **89.7%** | **1.91** | **[1.042,6.41]** |

## 4. Discussion

Several variables previously identified as critical in the MIA model were confirmed by applying penalized regression to our MIA dataset. Firstly, maternal weight loss and hypothermia were negatively associated with the MIA class confirming that the intensity of immune activation is critical as recently reported [10, 11]. Secondly, decreased litter size was associated with the MIA class, confirming that MIA could induce partial abortions and therefore reduce litter size as reported in some but not all studies [11]. Thirdly, higher BW was associated with the MIA class, a result which is in partial agreement with a study in which the fat mass was increased in the MIA adult progeny [25], but not with others in which MIA pups exhibited a decreased BW [26, 27]. Differences in dose or timing of poly(I:C) injection, as well as reduced sample size in previous studies could explain these discrepancies. Fourthly, decreased number of emitted USV emission at P8 was associated with the MIA class. USV are produced by pups in response to isolation from the mother, hunger and thermal changes, which call for maternal care: reduced number of USVs can therefore be interpreted as defects in the pup’s attachment behavior [15]. Our results are in line with studies showing that exposure to MIA induces USV communication deficits in young and adult MIA offspring [28, 29]. This has also been described in many ASD genetic models [15]. In contrast, Choi and his coworkers repeatedly found increased numbers of USVs produced by MIA pups [9, 30], a result that could be accounted for by different experimental conditions. Fifthly, in agreement with previous reports in adult offspring exposed to MIA induced by maternal influenza infection [31] or poly(I:C) injection [7], hypo-locomotion of pups (assessed by reduced distance travelled and time spent mobile) was associated with the MIA class. Altogether, these results indicate that penalized regression adequately selected variables previously identified as critical in the MIA model. In addition, penalized regression disclosed the possible involvement of four cytokines in MIA: CXCL10, IL-5, IL-15 and TNF-α. To our knowledge, IL-6 and IL-17A were the only cytokines previously connected to the neurobehavioral alterations in the MIA model, but only during the antenatal period [7-9]. Our work supports further exploration of the role of these four cytokines in the MIA model.

The data from this exploratory study should be replicated in other laboratories before being generalized. Other covariates such as the dose, timing and batch of poly(I:C), mouse strain, microbiota composition and behavioral variables collected in adults could be included therefore providing a more complete picture of the MIA model. We believe that our study not only provides valuable information on the characteristics of the MIA model in relation to the postnatal consequences of *in utero* exposure to MIA, but also highlight the importance of extending the use of MV statistics to consider confounding variables to explain any given result. Nevertheless, we wish to point that implementing penalized regression does not overcome the issue of a low sample size [32]. To summarize, this study illustrates the power of MV approaches, and the pertinence of penalized regression to analyze datasets from animal studies addressing complex inter-dependent phenotypes.

## Acknowledgements

We thank Lucien Relmy for technical assistance with animal care. We thank Cecil Czerkinsky for critical reading and editing of the manuscript.

